# Neural Correlates of a Trance Process and Alternative States of Consciousness in a Traditional Healer

**DOI:** 10.1101/2021.01.04.425271

**Authors:** Rebecca G. Rogerson, Rebecca Barnstaple, Joseph F.X. DeSouza

**Affiliations:** Centre for Vision Research, Interdisciplinary graduate studies, CAPnet VISTA, York University, M3P 1P3; Department of Dance and Neuroscience Graduate Diploma Program, CAPnet VISTA, York University, M3P 1P3; Department of Psychology, Biology, Canadian Action and Perception Network (CAPnet VISTA, York University, M3P 1P3

**Keywords:** brain, plasticity, trance processes, learning, auditory (A1), area prostriata, dance

## Abstract

Trance processes are a form of altered states of consciousness (ASC) widely reported across cultures. Entering these states is often linked to auditory stimuli such as singing, chanting, or rhythmic drumming. While scientific research into this phenomenon is relatively nascent, there is emerging interest in investigating the neural correlates of altered states of consciousness such as trance. This report aims to add to this field of ASC through exploring how the perception of an experienced *Sangoma*--traditional South African healer--entering a trance process correlates to BOLD signal modulation with auditory stimuli. Functional MRI data were analyzed using a General Linear model comparing music versus no music condition multiplied by the percept of experiencing trance (High or Low). Positive BOLD activation was shown in auditory cortex in both hemispheres during a trance process. Other brain regions tightly correlated to trance perception were right parietal, right frontal, and area prostriata (P<0.05, Bonferroni corrected). Orbitofrontal cortex (part of the Default Mode Network) was negatively activated and most correlated with music when trance was high, showing the largest differential between high and low trance perception. This is the first study to directly correlate BOLD signal variations in an expert subject’s percept of trance onset and intensity, providing insight into the neural signature and dynamics of this unique form of ASC. Future studies should examine in greater detail perception of trance processes in expert subjects, adding other neuroimaging modalities to further investigate how these brain regions are modulated by trance expertise.

## Introduction

Altered states of consciousness (ASC) have been described worldwide since time immemorial, often in association with religious or spiritual practices; until recently ASC were understood within biomedical paradigms as a form of psychopathology or only as a physiologlcal response to stress (Boddy, 1994). While there is growing interest in conducting experimental investigations of ASC such as those associated with hypnosis, meditation, flow states, or disassociation the neurological/biological origins or effects of trance are still largely unknown (Flor-Henry, Shapiro, Sombrun, & Walla, 2017; Hove et al., 2016; Peres et al., 2012).

Trance processes can vary across ethnicities and socio-cultural practices, as well as within traditions, communities, and even among individuals. Therefore, a single definition and understanding of trance cannot be reduced, naturalized, medicalized or universalized, but is best recognized “on its own unique terms” (Boddy, 1994). What can be said, however, is that trance processes commonly display “alterations or discontinuity in consciousness, awareness, personality, or other aspects of physical functioning” (Cohen, 2007). Another feature that trance processes share is that they are highly absorptive and may resemble or induce the experience of *flow*, which Csikszentmihalyi defines as an *optimal experience* in which skills and action are perfectly matched, resulting in a “forgetting of all else” (Seligman, & Kirmayer, 2008). Lastly, trance processes are often facilitated or accompanied by auditory stimuli such as chanting, singing, or rhythmic drumming. In the *Ngoma* traditional healing practices of South Africa – the tradition which the participant practices - the *thwasa*, or healing student, learns throughout the community-based apprenticeship process how to “call up the ancestors” through a trance process with the use of these internal and external auditory stimuli and enters into respective flow states. For this study, the music was selected by the subject to induce trance within the confines of the MRI machine, with the goal of examining the experience or *perception* of the trance process. The expert subject has over 20 years of experience with trance processes as a practicing *Sangoma*. Our approach to this study is scaffolded on simultaneous research involving groups of experts (break-dancers, ballet dancers, choreographers) who regularly engage in specific movement/performance behaviors as part of their work environment. In one of these investigations (Olshansky et al, 2014), we had an expert break-dancer from New York select a 1-minute piece of music that he used for many years in break dancing battles, and with which he was highly familiar over his 20-years of dancing. Experts have putative specialized neural circuits for behaviors related to their area of expertise that can be probed for brain-behavior relationships (Olshansky, Bar, Fogarty, & DeSouza, 2014) and can be trained to reduce neurodegeneration((Zilidou et al., 2018; Douka, Zilidou, Lilou, & Tsolaki, 2019)). Specifically, in this case study with a *Sangoma*, our aim was to examine brain regions showing an interaction with the expert subjects’ percepts, correlating increases in modulation with a high/low perception of trance. We used functionally mapped brain regions of interest from Hove et al., (2016) and Bar & DeSouza (2016) to anatomically guide our inquiry in trance perception.

## Materials and Methods

### Participants

One experienced practicing *Sangoma* was scanned (female, age 42 years, dance experience of +20yrs). York University’s ethics committee approved the study (e2013-313), and written informed consent was obtained from the participant in accordance with the committee’s guidelines.

### Equipment and Scanning Procedure

A 3T Siemens Tim Trio MRI scanner was used to acquire functional and anatomical images using a 32-channel head coil. T2*-weighted echo planar imaging using parallel imaging (GRAPPA) with an acceleration factor of 2X with the following parameters: 32 slices, 56 × 70 matrix, 210 mm × 168 mm FOV, 3 × 3 x 4 mm slice thick, TE = 30 ms, flip angle of 90°, volume acquisition time of 2.0 s, was used. There was a total of 240 volumes per scan. Echo-planar images were co-registered with the high-resolution (1 mm^3^) anatomical scan of the subject’s brain taken at the end of the session (spin echo, TR = 1900 ms, TE = 2.52 ms, flip angle = 9°, 256 × 256 matrix). The subject’s head was restrained with padded cushions to reduce movements. While in the scanner, our expert wore headphones to hear the music. The trance inducing music task employed a blocked design 30 seconds *OFF* and 60 seconds *ON. ON* states were alternated five times. These tasks were analyzed using the General Linear Model (GLM) in BrainVoyager QX (Brain Innovation v2.1.1.1542, Maastricht, The Netherlands) with the boxcar function convolved with a double gamma hemodynamic response function and her perception of trance. Following statistical analysis of the BOLD signal data was conducted in MATLAB (The MathWorks Inc., version 8.4.0.150421, 2014b).

### Task Procedure

While in the scanner there were two tasks: (1) a trance music-inducing task cued by music and (2) moving the right foot at 1-Hz (Olshansky, Bar, Fogarty, & DeSouza, 2014). During the trance task, the subject was told to listen to the music^1^ and attempt to enter into trance as she would in her practice as a *Sangoma*. Immediately following the completion of the scan, while the subject was still in the scanner, we asked the participant to rate the success of each of the five 1-minute pieces of music and whether she achieved a trance state (1-High) or not (0-Low). A follow-up interview was conducted approximately 20-minutes later, after the structural scans were completed and the subject left the MRI suite.

### Preprocessing

Functional data were superimposed on anatomical brain images, aligned on the anterior commissure-posterior commissure (AC-PC) line, and transformed into Talairach space. Functional data from each scan was screened for motion artifacts from head movement or magnet artifacts to detect eventual abrupt movements of the head. In addition, we ensured that no obvious motion artifacts were present in the activation maps.

### Statistical Analysis

The subject’s functional data was analyzed using the general linear model (GLM) module with a weighting of a perceptual model convolved with the music block compared to the period where there was no music (fixation blocks-light grey periods from Figure 1 BOLD signal). Bonferroni corrections was used (P<0.0001) since it is more conservative than False discovery rate (FDR). Data from GLM regions were extracted and pair-wise *t*-tests were used to compare BOLD signal differences from brain regions (Auditory Cortex (left and right hemispheres), Supplementary Motor Area (SMA), Parietal activations (PPC), visual cortex, area prostriata, anterior cingulate cortex (ACC) and posterior cingulate (PCC), motor cortex area of the right foot, caudate and putamen from Basal ganglia. The last three brain areas were derived from Bar and DeSouza (2016) locations.

**Figure 1:**
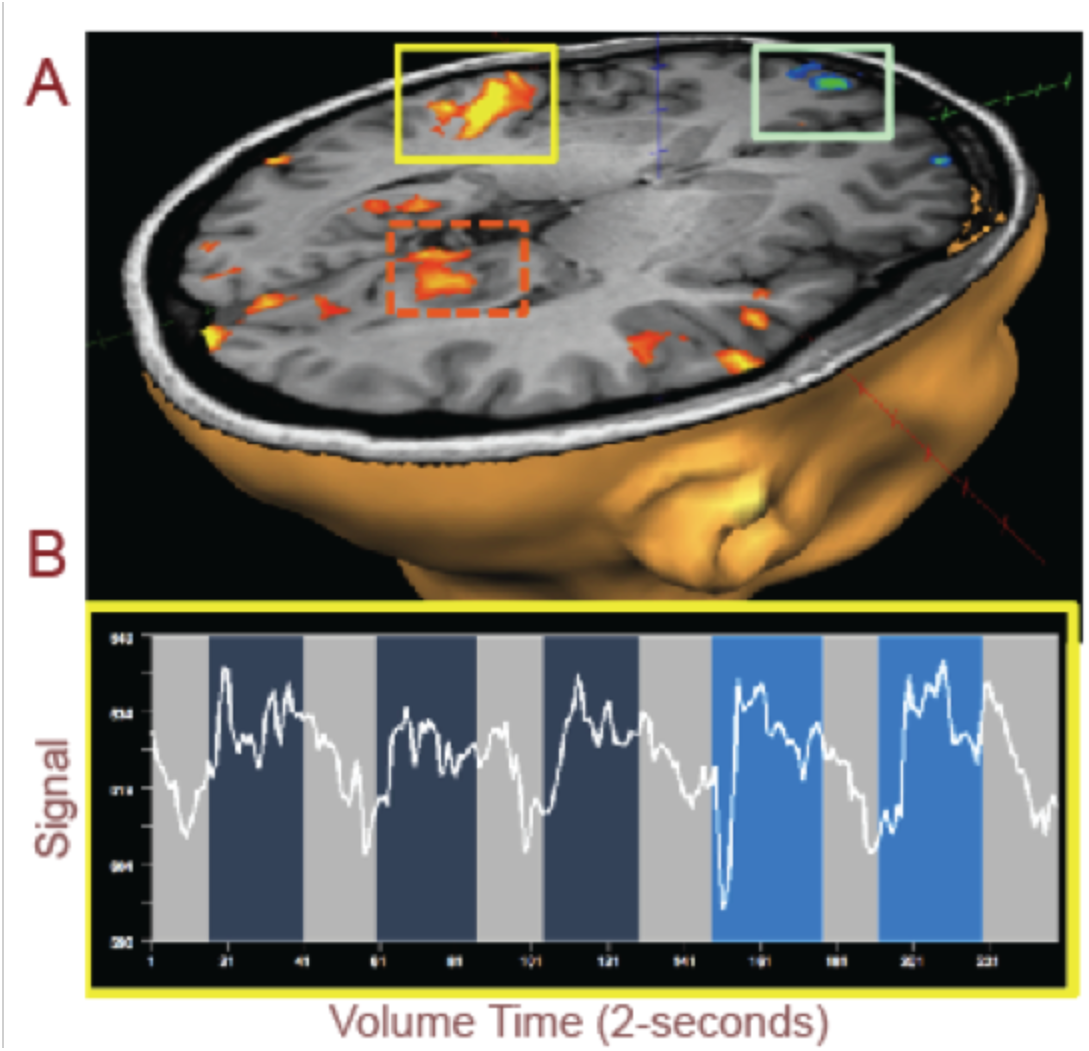
A) Auditory cortex (yellow box) shows an increase in BOLD signals while listening to the music to induce trance state compared to no music. The light green box surrounds orbitofrontal cortex. The orange dashed box highlights area prostriata. B) BOLD signals extracted from the auditory cortex (yellow box above) showing increased signals for music with the last two blocks showing an increase in trance perception.

## Results

The subject rated two blocks achieving the trance state (High) during her self-selected music. This rate of achieving trance in the confines of the MRI scanner was expected since the subject was asked to be immobile and not move. The GLM contrasting perceived trance induced by preferred music was contrasted to fixation showed brain regions within auditory cortex, area prostriata, visual, parietal cortex and orbitofrontal (P<0.05, Bonferroni corrected; threshold of k=8 voxels; Forman et al., 1995). Both hemispheres within auditory cortex within the temporal lobe showed an increase in BOLD signals when perceived trance was high; Fig 1A (yellow box) & 1B highlight one hemisphere auditory cortex and the BOLD signal from the fMRI scan. The BOLD signal (white line Fig 1B) starts out low (light grey highlighted blocks of time), then increases in amplitude soon after the music begins (darker blue); this lasts for a 1-minute time period, after which the BOLD signal reduces when there is no music. This pattern is repeated four more times. Brighter blue depicts the blocks of music when the subject perceived a higher perception of trance (rated as 1). When we average across perceptual blocks of trance from this auditory region and then parse the subject’s “low” or “high” perception of trance, we observe that there is a significantly higher BOLD signal than for low perception (Fig 2). The exact same auditory stimulus was played for each 1-minute block. When we average across the blocks of 1-minute of music for the dark blue and brighter blue blocked regions in Fig 1B, we produce Fig 2A with blue and grey lines for periods with music and white lines for periods with no music. The bar graphs in Fig 2B show times at which the perception of trance was significantly higher than when the same music was perceived as a low trance (paired *t*-test, P<0.05), thus it was the subject’s *perception* of trance that putatively modulated the auditory signal input since the input to the headphones was identical each time.

**Figure 2:**
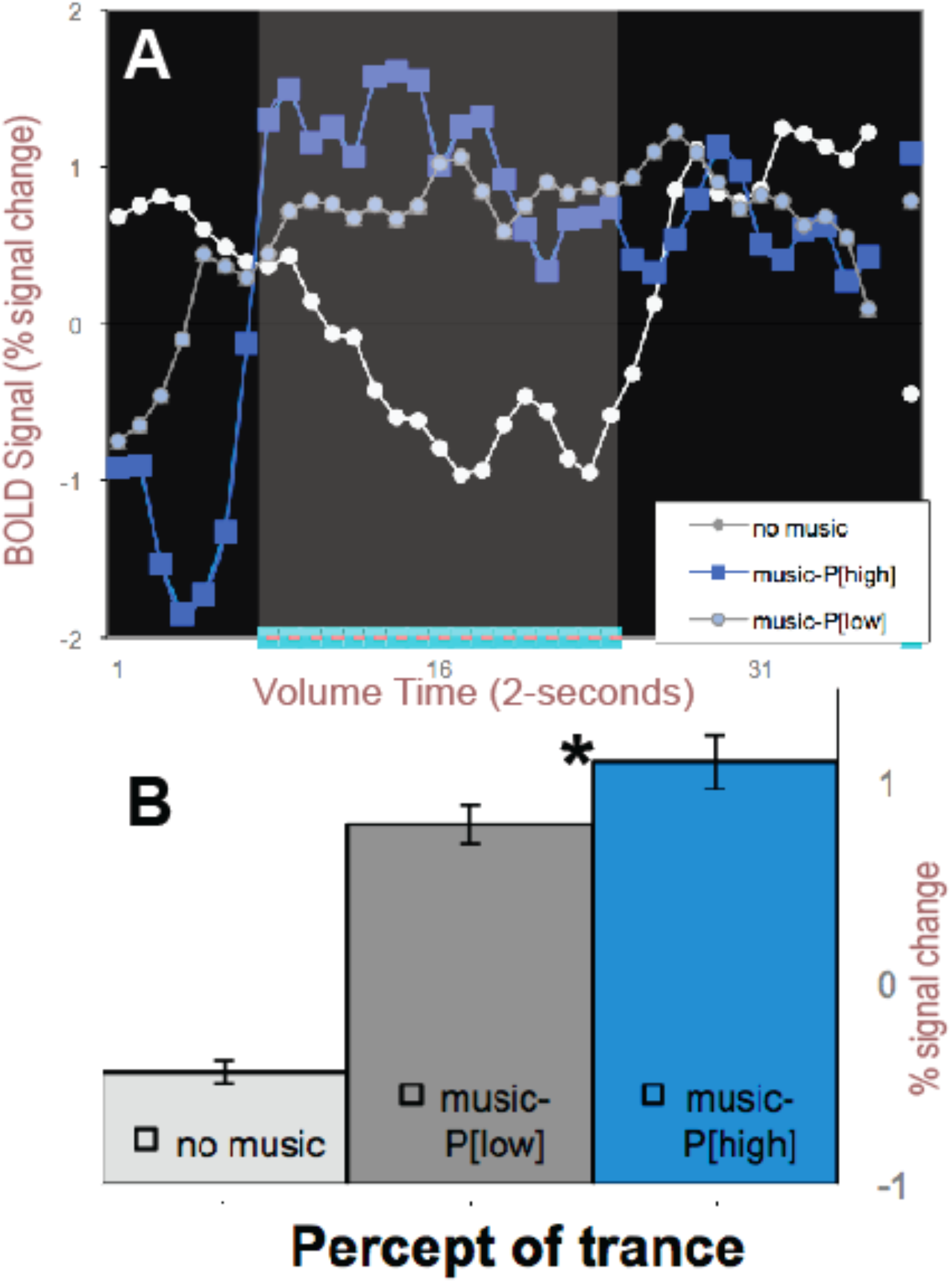
Right Auditory cortex BOLD signals averaged across perceptual states of High and Low trance percept. A) BOLD signal averaged across the 5-blocks of 1-minute of music and color-coded (blue=high; grey=low) perception of trance. White line depicts when no music was played over the headphones in the MRI suite. B) average BOLD signal for the three types of data. * -signifies P<0.01 paired *t*-test.

We now plot this right auditory cortex region comparing perception of trance in the Low (*P[LOW]*) and High states (*P[HIGH]*) to compare it with all other regions from the GLM analysis that passed our spatial Bonferroni threshold and spatial voxel clustering (Forman et al., 1995) in Fig 3. But first we need to discuss the format of the following summary figure which includes all the brain regions that were mapped with the GLM and from anatomical locations pulled from the Hove et al. (2016) and Bar and DeSouza (2016) studies.

**Figure 3.**
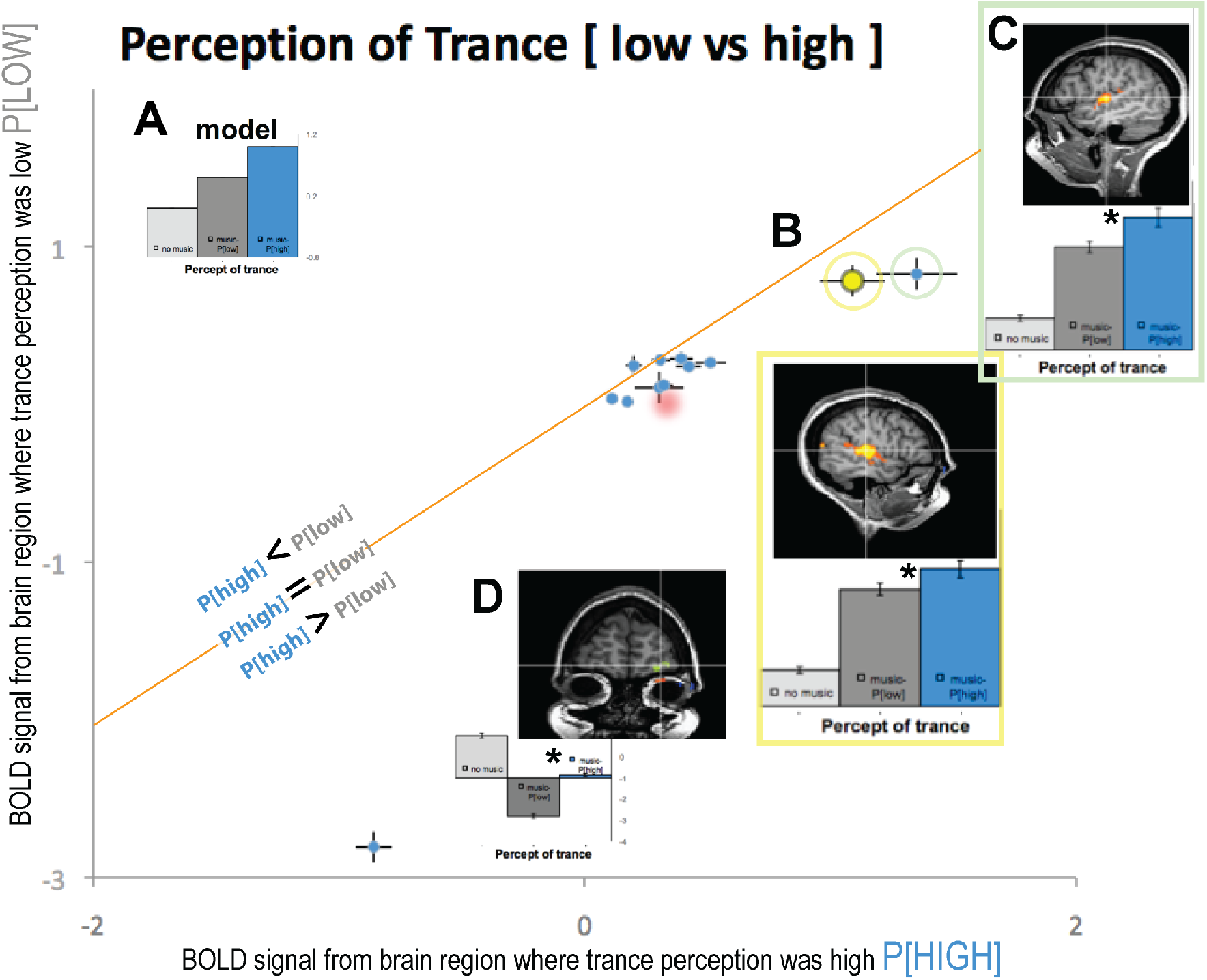
A) The Model BOLD signal if trance has a P[*HIGH*] modulated by the music signal. Diagonal orange line represents where low and high trance perception is equal with data below and above the line as the differences. B) Same brain area as in Fig 1A from right auditory cortex. C) left auditory cortex bar plots and location. D) Orbitofrontal cortex bar plot and spatial location. Other data points that were examined from functional ROIs clustered near orange line. Red drop shading for two data points signify anatomical regions from Hove et al., 2016 study. * - denotes significant P<0.05

Figure 3A inset bar graphs is for an idealized model BOLD signal that would be observed for a brain region significantly modulated by the perception of trance. Plotting the ratio of high to low trance perception [*P*], the orange diagonal line in the large figure represents where trance signals would be equivalent (*P[high]* = *P[low]*) for a specific brain region. Three brain regions, early right hemisphere visual cortex, left parietal and cerebellum activations both fall near this diagonal line and are thus *NOT* significantly modulated by perception of trance (P’s>0.05). In our results, all other data points fall to the right or below this diagonal orange line, indicating a stronger signal for HIGH perception of trance compared to LOW perception (P<0.05). If these were to the left of the diagonal line, that would have indicated a higher signal for the LOW perception of trance (*P[low]*) than the HIGH (*P[high]*) with no functionally mapped brain region was significantly to the left of the line. Only the right visual cortex data point is to the left, but it is not significantly far enough to fit this model.

We plotted the auditory cortex signal from Fig 1A and Fig 2B as data point B in Figure 3 (yellow data plot) which is below the diagonal line, illustrating that there is a significant difference between HIGH and LOW perception of trance (P<0.05). The error bars of this Fig 3B data point plot standard error of the mean signal across high and low perceptual states, and these error bars do not touch the orange diagonal line.

Additionally, left auditory cortex showed an increased BOLD signal for trance perception (green box, Fig 3C) which was stronger than that from the right hemisphere. Area prostriata, right parietal, foot motor area^Bar+DeSouza^, caudate^Bar+DeSouza^, putamen^Bar+DeSouza^ and supplementary motor cortex^Bar+DeSouza^ also showed stronger activation (Fig 3). These brain regions showed significantly enhanced signal for high trance perception, with the exception of right visual cortex, left parietal and cerebellum, whose data points touched the line of unity. We used anatomical locations of PPC^Hove^, ACC^Hove^ from Hove et al., (2016) to further probe our data (red drop shaded area in Fig 3) and found that these areas were also associated with a higher percept for trance (P<0.05).

Most interestingly, the orbitofrontal cortex (OFC), an area activated in the default mode network (DMN) when a task is stopped and the subject is at rest, was most negatively correlated to the perception of trance and showed the largest difference of high *(P[high])* compared to low (*P[low]*) trance perception (Fig 3D). OFC was the furthest data point from the diagonal line, indicating that it was the brain area that showed the largest difference for trance perception in our subject (*P[high]*>>*P[low]*). When we examine the BOLD signals from OFC, there is an expected a modulation in the DMN up to the point in which the subject’s perception of trance increased to *high*, at which point the DMN showed decreased activation for the remaining period of data collection (see supplementary figure S1). To our knowledge, this could be the BOLD signal neural correlate of trance (ASC) beginning since the DMN is turned off and this OFC BOLD signal does not modulate down when music is back on. This may be an indication that culturally specific auditory stimuli are involved in triggering a trance process in the participant, or it could demonstrate her experience or familiarity with auditory stimuli used while learning trance processes during her apprenticeship in South Africa in 1997.

## Discussion

Trance processes represent an intriguing form of ASC with aspects of volitional control not unlike meditation and hypnosis (Hove et al., 2016). Of particular interest for neuroscience is the correlation of the associated experience or perception of the altered state with brain imaging data. Research of this phenomenon may be a rich source of information as to how attention, volition, activation, and/or suppression of neural networks can play a role in consciousness (DeSouza, & Everling, 2004; DeSouza, Menon, & Everling, 2003; DeSouza, Ovaysikia, & Pynn, 2012). In 2015, The Martinos Centre and Max Planck Institute conducted research with fifteen self-identifying German and Austrian shamans to probe brain network reconfiguration and perceptual decoupling during trance (Williams, 2015; Rogerson, 2017). Additionally, using trance-inducing music, Hove et al., (2016) found that the effects of drumming on sensory input created patterns of network engagement that they suggest are involved in trance. This included areas in posterior cingulate cortex (PCC), dorsal anterior cingulate cortex (dACC), and left insula/operculum (Hove et al., 2016). They also found increased coactivation of the PCC with the dACC and insula, key hubs in the DMN and executive control networks. Decoupling of seeds within the auditory areas might indicate suppression of the auditory stimuli during trance, with this suppression seeming to play a key role in entering an ASC (Hove et al., 2016). Through seed-based functional connectivity and a second seed-based analysis they found a higher degree of functional connectivity from PPC to ACC/insula, along with clustered activity in the caudal pons, while larger-scale network connectivity was also enhanced during trance. Their seeded regions in auditory cortex suggest it is less connected to PPC and ACC/insula regions. They concluded that trance involves more sustained task maintenance and cooperation of brain networks associated with internal thoughts and cognitive controls, coactive defaults, control networks, and decoupled sensory processing than once believed (Hove et al., 2016; Rogerson, 2017).

Using regions from Hove et al. (2016) in our subject as anatomical localizers to probe whether these brain regions are modulated by trance in our subject, we found that they (PPC^Hove^ and ACC^Hove^ regions) were indeed an increased modulation by HIGH perception of trance, but our functionally mapped regions in auditory and OFC regions showed much larger modulation by HIGH perception of trance. In a departure from their study, we did not rely on data modeling to infer higher trance-related signals in the seed regions; we took a different approach by asking the subject when trance perception was high and low and relied on behavior as a model to run our analysis. While Hove et al (2016) included self-reports following the trance condition to probe whether subjects felt they had entered a trance process, they did not include more nuanced measures of alterations in the degree of trance during this condition. We believe probing the subject’s perceptual shifts during an enhanced trance process adds to their findings and should be part of standard operating procedures to model future trance processes in examinations of ASC studies of trance perception. Our experiment is the first to show increased BOLD signals correlated with the participant’s perception of trance in brain regions activated through music to induce a trance process in an experienced *Sangoma*.

While there is limited neuroscientific research specifically on trance processes, there is growing interest in the investigation of other kinds of ASC that may suggest future directions for inquiry (Crick, & Koch, 1990). The Stanford University School of Medicine conducted a hypnosis-focused study in 2016 with 57 participants, finding that decreased activity in the dorsal anterior cingulate and increased activity in the dorsolateral prefrontal cortex and the insula occurred during a hypnotic state. Blanke’s work on Out of Body Experiences (OBE) which he purports are experienced by 2-5% of the population, shows that brain function disruptions also occur for people who experience OBE. A splitting or “body as the observer” mode is experienced, which may be the result of enhanced activity in the parietal zone area (Rogerson, 2017). Coactivation may also play a role with fluctuations in levels of consciousness, awareness and sensory patterning (or lack thereof) occurring (Williams, 2015; Rogerson, 2017; Samuel, & Bettina, 2010). The right parietal region in our study’s participant is near Blanke’s OBE regions. Her right parietal brain regions were modulated by high and low trance, but the left hemisphere region was not modulated by trance. It is interesting that the right parietal in our study shows modulation by trance and the OBE is typically right hemisphere. The right parietal region may therefore be important for future studies interested in ASC.

In Csikszentmihalyi’*s Flow: The Psychology of Optimal Experience*, he argues that regular ‘practicing’ of, and engagement in, absorptive activities may result in *a flow state*, defined as an automatic, effortless and highly focused state of mind that occurs when self-reflexive consciousness is harmoniously ordered (2009). Through *optimal experience* a sense of exhilaration and deep enjoyment occurs creating a landmark in the memory for what life should look like (Rogerson, 2017; Csikszentmihalyi, 2009).

The association of trance processes with music suggests the involvement of auditory cortex and areas previously described in a group analysis (Hove et al., 2016), but what has not been previously described or investigated are brain regions associated with self-referential memories and experiences that may then trigger trance processes. There is a need for studies examining regions involved in memory and emotion coding to investigate whether these may suppress or encourage trance processes. With regards to the latter statement, the area prostriata brain region is situated near input areas of visual and auditory cortex (Rogerson et al., 2018 IMRF abstract); more importantly, it has strong interconnections with the limbic and retrosplenial cortex in anatomical tracing studies ((Cavada, Compañy, Tejedor, Cruz-Rizzolo, & Reinoso-Suárez, 2000; Rosa, Casagrande, Preuss, & Kaas, 1997; Rosa, Soares, Fiorani, & Gattass, 1993; Rosier, Leroux, Vaudry, Orban, & Vandesande, 1991; Sousa, Piñon, Gattass, & Rosa, 1991). Area prostriata is anterior to the visual cortex and has been described as a visual area that acts as an interface for peripheral vision and fast processing (Tamietto, & Leopold, 2018), and may also be involved in sending information to multisensory and higher order association areas in temporal, cingulate and orbitofrontal cortex (Rockland, 2012; Yu, Chaplin, Davies, Verma, & Rosa, 2012). Our study is the first to uncover this putative functional connection as we did not use visual stimuli to functionally activate area prostriata, but rather music associated with the subject’s ongoing trance processes. Thus, this could be putatively from her “a priori” experiences with trance from memory areas that “turn on” area prostriata when music stimuli are present for trance.

### Limitations

One limitation is that our current study entailed only one fMRI scan of 5-minutes of music and 3-minutes of non-music. More data is required with this participant as well as other healers who practice trance processes, using the exact same scan sequence and MRI as used in the previous case study (Olshansky et al., 2014). In consideration of this, we used a very conservative threshold of Bonferroni correction and clustering (Forman et al. 1995) to ensure our regions could be discussed in this case study and to avoid jumping beyond what the case study model allows us. Therefore, future studies will seek to confirm the modulation of trance processes in regions of the default mode network such as orbitofrontal cortex, and whether area prostriata together with other regions in this case study can be replicated for a group of *Zangoma*.

## Conclusion

We have aimed to contribute to a reliable neural marker of trance processes (Jureidini, 2004) in a case study of an experienced *Sangoma*. We have demonstrated an effect of a heightened trance process with associated brain regions that were modulated by her perception of trance. This perceived correlation was within the “ON-music-state” bilaterally in auditory cortexes, visual/parietal areas, and most interestingly, modulated over into the default “OFF-state” only within bilateral orbitofrontal cortex. Since it was only a single case study, we were very conservative in describing the brain regions that were activated, using the most conservative threshold of the Bonferroni multiple comparisons and also used a next neighbor threshold that increased our confidence and reduced our false positive chances. Our future research aims to examine in greater detail trance percepts in this subject and in other similarly skilled participants. We feel that these kinds of interdisciplinary and collaborative research endeavors may be beneficial to establishing new kinds of clinical approaches and ongoing research projects (Seligman, & Kirmayer, 2008).

## ACKNOWLEDGEMENTS

A National Science and Engineering Research Council (NSERC) Discovery Grant (2017-05647) to JFXD funded this project.

---

**Figure S1:**
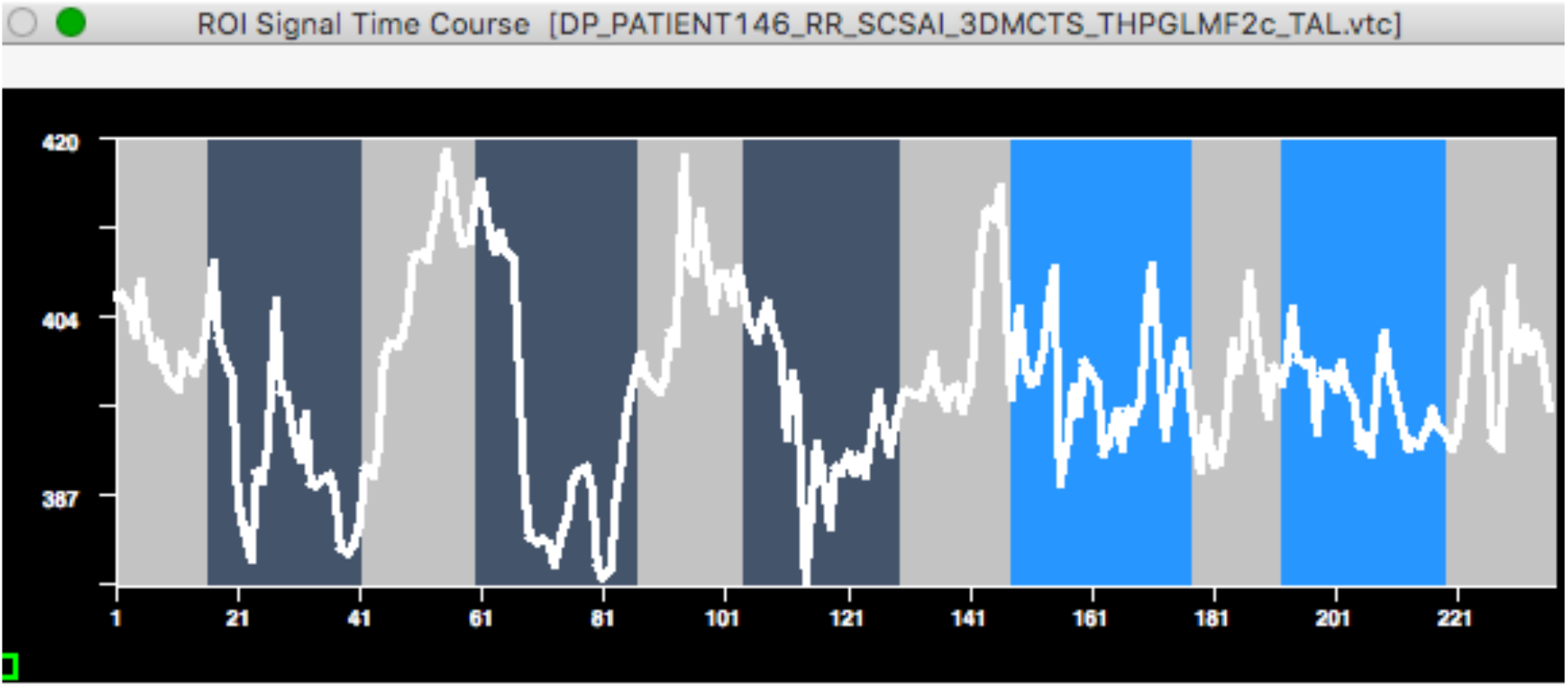
Orbitofrontal cortex (OFC) brain activation showing modulation of the Default Mode Network at Volume time (141) and then when the subject’s perception of trance was HIGH this OFC regions than has a change from DMN activity to being shut off from Volume time (141 to end). This suggests an interaction between the DMN and trance with her perception of trance.

## Footnotes

1 The music stimulus used was a one-minute excerpt on YouTube, taken during Swaziland Cultural Group performance. https://www.youtube.com/watch?v=LJGyAxYfojk

